# Anatomically informed multi-level fiber tractography

**DOI:** 10.1101/2020.12.16.423042

**Authors:** Andrey Zhylka, Alexander Leemans, Josien Pluim, Alberto De Luca

## Abstract

Diffusion weighted MR imaging can assist preoperative planning by reconstructing the trajectory of eloquent fiber pathways. A common task is the delineation of the corticospinal tract in its full extent because lesions to this bundle can severely affect the quality of life. However, this is challenging as existing tractography algorithms typically produce either incomplete results or multiple false-positive tracts. In this work, we suggest a novel approach to fiber tractography that reconstructs multi-level structures by progressively taking into account previously unused fiber orientations. Anatomical priors are used in order to minimize the number of false-positive pathways. The devised method was evaluated on synthetic data with different noise levels. Additionally, it was tested on in-vivo data by reconstructing the corticospinal tract and it was compared to conventional deterministic and probabilistic approaches. The corticospinal tract reconstructed by our method includes lateral projections that could not be observed with deterministic methods, while avoiding spurious tracts reconstructed by probabilistic tractography. Furthermore, the proposed algorithm preserves the neuroanatomical topology of the pathways to a larger extent as compared to probabilistic tractography.

## 1 Introduction

Diffusion MRI provides an opportunity to estimate fiber orientations through the brownian motion of water molecules. This imaging technique allows for exploring brain connectivity in-vivo and non-invasively (Fornito et al.,2013;Hagmann et al., 2007) as well as performing virtual dissection (Jacquesson et al.,2018;Panesar et al., 2019;Thiebaut de Schotten et al.,2011;Catani & Thiebaut de Schotten,2008), aiding presurgical planning (Jones,2010) and serving as a reference during surgery (Costabile et al.,2019). Despite its promising results, fiber tractography remains challenging, as the results of existing methods have been shown to perform satisfactory on either sensitivity or specificity, but not both (Maier-Hein et al.,2017;Schilling et al.,2019; Sarwar et al.,2019).

One of the origins of the tractography shortcomings is that the brain fiber network contains multiple crossings and other complex co-occurrences of fiber bundles, which are hard to resolve (Behrens et al.,2007;Jeurissen et al.,2013). Multiple diffusion models have been proposed to reconstruct the organization of fibers from the diffusion signal, e.g. the diffusion tensor (DT) model (Basser et al.,1994), diffusion kurtosis imaging (Jensen et al.,2005) and the estimation of the fiber orientation distribution (FOD) with spherical deconvolution techniques (Tournier et al.,2007). Based on the way tractography methods use the information provided by such models, they can be categorized as deterministic or probabilistic. Deterministic approaches follow either the dominant diffusion (or fiber) direction (Mori et al.,1999) or one of the main directions that is the least deviating from the orientation of a previous step (Tournier et al., 2007). On the other hand, probabilistic approaches sample and propagate orientations based on the FOD in the voxel (Tournier et al.,2012). Even though probabilistic methods are able to reconstruct most of the true-positive pathways, they tend to have a high false positive rate (Maier-Hein et al.,2017). In contrast, deterministic methods are prone to generating false negative results, but their results are reproducible by definition.

Apart from the intrinsic flaws of existing tractography algorithms, the results of fiber reconstruction are affected by the constraints that are imposed to achieve anatomically plausible results. Thus, branching pathways are quite likely to be pruned due to the angular constraints and spatial resolution, leading to an increased false negative rate (Mortazavi et al.,2018).

For the purposes of surgery planning and virtual dissection, the sensitivity of tractography plays a key role, as the correct prediction of the extent of resection is essential to avoid functional impairment. The corticospinal tract (CST) is one of the bundles which neurosurgeons and neuroradiologists focus on during surgery planning to prevent motor function degradation (Weiss et al.,2015). Despite the existence of guidelines to reconstruct the CST (Catani & Thiebaut de Schotten,2008;Wakana et al.,2007), achieving a complete dissection of this bundle is challenging, due to fibers passing through the centrum semiovale, which often leads to underrepresented reconstruction of the face-related part of the motor cortex (Fortin et al.,2012). Additionally, for longitudinal studies involving patients with brain injuries it is important to maintain the organizational patterns typical of the CST, their topographic organization (Lee et al.,2016). Preserving such feature is likely challenging also for probabilistic approaches unless it is specifically taken into account (Aydogan & Shi,2018).

Given that anatomical landmarks are well defined for the CST (Thiebaut de Schotten et al.,2011;Wassermann et al.,2016;Han et al.,2010), incorporating anatomical prior knowledge in the tractography might offer a viable solution to improve its performance. For instance, the bundle-specific approach MAGNET (Chamberland et al., 2017) enforces a specific direction for tract propagation by accepting a user-defined regions of interest (ROI) to enhance the reconstruction of the optical pathways. A similar guidance of the fiber tracking can be achieved also using transcranial magnetic stimulation to find the brain regions responsible for specific functionality for the purpose of filtering fiber bundles related to those regions (Negwer et al.,2017). The anatomy-driven automated approach, TRACULA (Yendiki et al.,2011), generates a distribution of fiber bundles based on a manually delineated fiber atlas. However, in TRACULA the atlas is constructed using FACT (Mori et al.,1999), which, by definition, has limited performance in case of complex fiber configurations. A more flexible way of including anatomical information was developed recently in the bundle-specific tractography framework (Rheault et al.,2019). It uses prior knowledge about certain fiber bundles to enhance the FOD so that they are closer to representing orientation distributions of the target bundles. This, in turn, improves the tracking results for the target bundles. The main drawback of this method is its strong dependence on a template. If the anatomy associated with the template is not representative, similar limitations will be present in the results of the BST algorithm.

Most anatomy-aware approaches attempt thus to either improve the streamline propagation or to enhance the FOD estimation. However, the aforementioned methods do not exploit all information available in the FOD. For one, the possibility of incorporating fiber bifurcations with high angular deviations along fiber trajectories is not taken into account in most of the existing approaches. This problem has been first investigated in Descoteaux et al.(2009) by introducing the concept of pathway splitting, but the proposed framework is prone to a high false positive rate due to a lack of control over the products of the splitting procedure. Additionally, extensive splitting at every point is not efficient as it not only leads to expanding the target bundle, but also captures intersecting bundles and reconstructs them.

In this work, we propose a novel approach to fiber tractography that adds branches to the pathways that have been previously reconstructed but do not reach a predefined target region. Repeating this procedure, each time we generate a new set of branches. Each set aims to improve the extent of the bundle and represents a new bundle level. This way the algorithm iteratively improves the fiber reconstruction, resulting in a multi-level structure (Fig. 1). By defining target and seed regions based on anatomical priors, the algorithm imposes constraints on the reconstructed streamlines, limiting the number of false-positive reconstructions. This concept can be fused with a wide range of tractography algorithms, e.g. any algorithm relying on an FOD. In this work we focus on the proposed multi-level strategy in combination with deterministic constrained spherical deconvolution (CSD) based tractography (Jeurissen et al.,2011).

**Figure 1:**
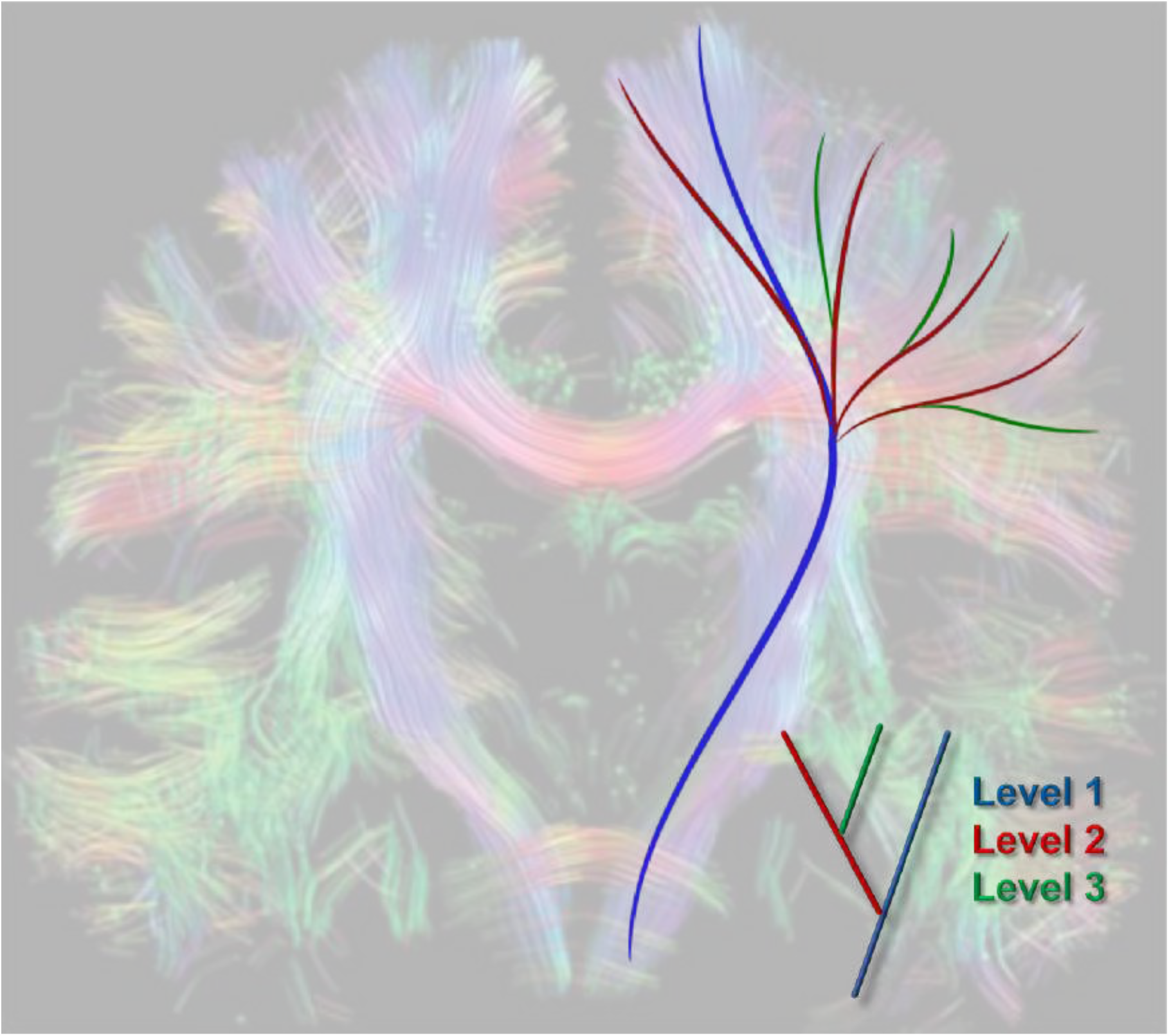
Current fiber tractography methods such as deterministic FOD-based tractography reconstruct only a subset of the pathways (blue). However, by propagating along the FOD orientations that were not used by a conventional tractography algorithm, the reconstruction can be iteratively extended by adding new sets of branches per iteration (red and green). This leads to a final tractography result consisting of multiple levels. The background picture of the whole-brain fiber tractography result is taken from Tournier et al. (2011) with permission.

We demonstrate that our algorithm combines the best of two worlds (probabilistic and deterministic tractogrpahy) by providing robust, reproducible and locally coherent pathways with an adequate spatial extent and a preserved topographic organization.

## 2 Material and methods

### 2.1 Algorithm

The core of our algorithm is a novel multi-level fiber tractography (MLFT) strategy that is applicable to a wide range of fiber tractography methods to take potential pathway splittings into account. It is an iterative procedure that requires user-defined starting and target regions as well as stopping criteria. Without loss of generality, we choose to focus on combining our tractography strategy with the deterministic CSD-based streamline tracking (Jeurissen et al., 2011).

Our algorithm iteratively expands the results with new branches. The set of branches added at one iteration can be considered one level of the overall bundle reconstruction. Firstly, at each iteration conventional deterministic CSD-based tractography is performed. Secondly, every point of the pathways reconstructed at the preceding level that corresponds to multiple FOD peaks is added to the new set of seed points, provided that those pathways do not enter the target region. Initial directions are defined as the FOD peaks of the new seed points that were ignored during the reconstruction of the preceding level. The algorithm runs for a pre-defined number of iterations or until a pre-defined convergence criterion is met as, for instance, coverage of the target area. Finally, the tracts that do not enter the target region are discarded (Fig. 2).

**Figure 2:**
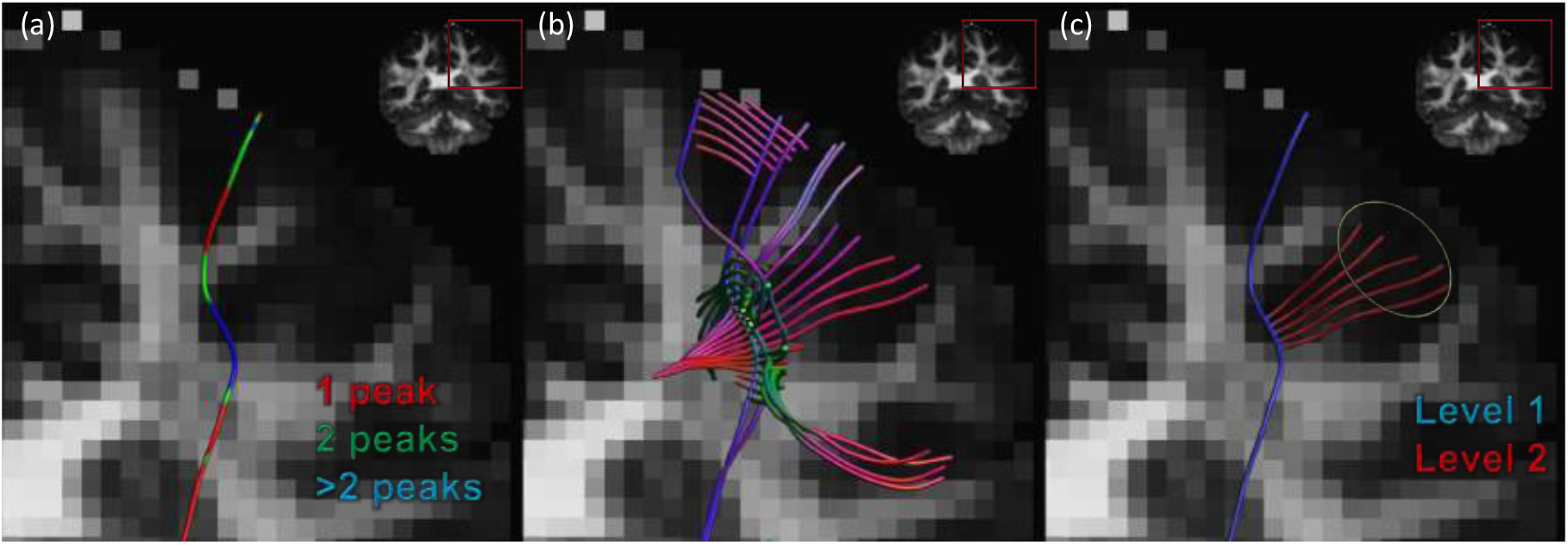
The pipeline of the algorithm. (a) The tract produced by the deterministic CSD-based tractography includes points with multiple FOD peaks, some of which are ignored. (b) Using these points as seeds with the unused peaks as initial locations, another iteration of CSD-based tracking is performed to obtain a new level of the result. (c) In the last stage only the tracts that enter the predefined target region are retained.

### 2.2 Data

We performed experiments on both simulated and acquired diffusion weighted images. A numeric phantom was generated using *ExploreDTI* (Leemans et al., 2009) with 6 volumes at b = 0 s/mm^2^ and 60 volumes at b = 1200 s/mm^2^ with a resolution of 1 mm isotropic (Fig. 3). The phantom represented three fibers with high overlap and two branching spots and was inspired by physiologically plausible fiber configurations as those that can be observed in the CST. The experiments with this phantom were performed without noise and for two SNR levels: 25 and 15.

**Figure 3:**
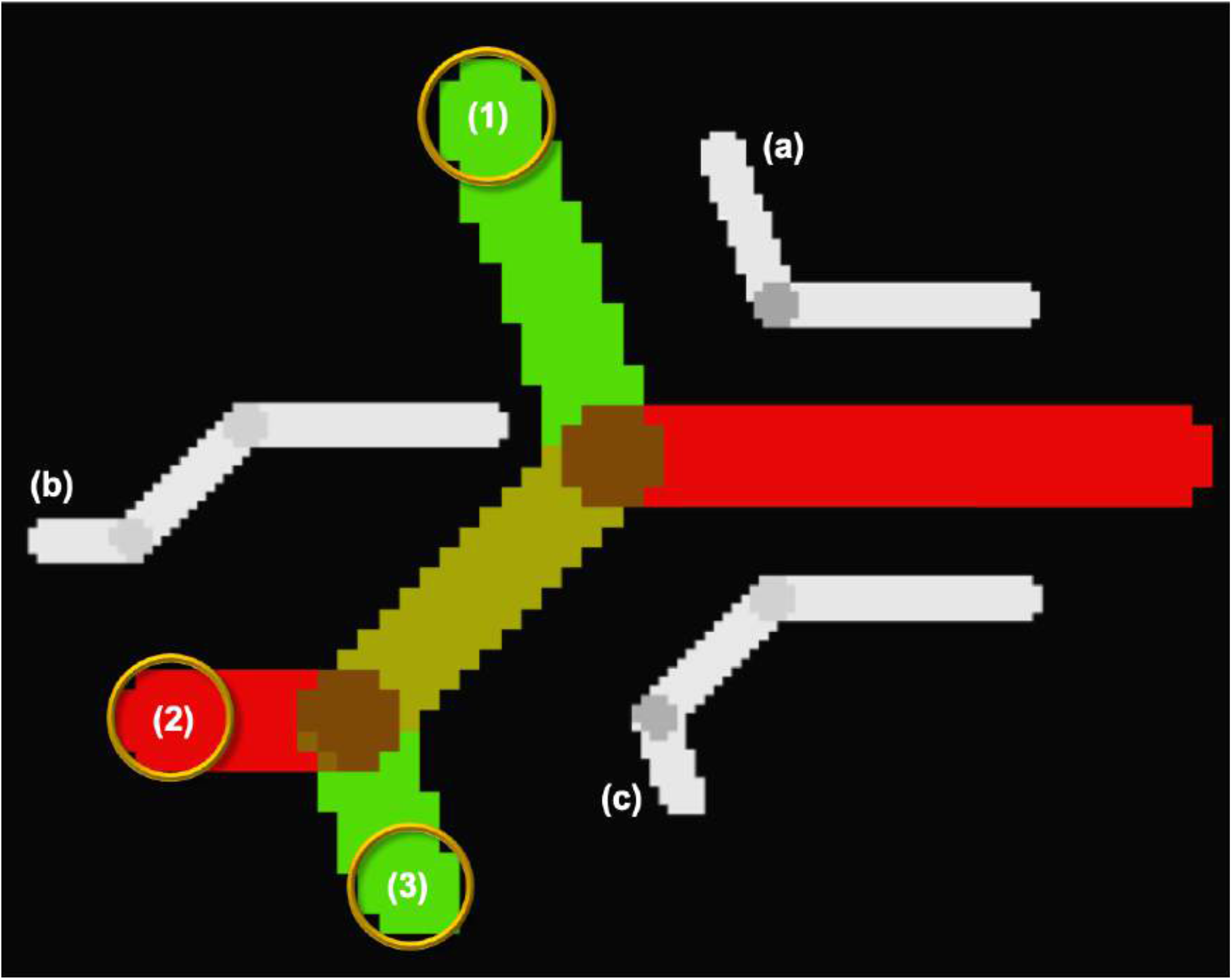
A representation of the numeric phantom (colored based on diffusion direction and fractional anisotropy (FA) value) with two branching points. It consists of three individual fibers (a-c, colored according to FA value) with corresponding endpoint regions (1-3).

In order to analyze the performance of our method on in-vivo brain images, the *MASSIVE* (Froeling et al., 2017) dataset was used. The data consisted of 430 volumes at b = 0 s/mm^2^, 250 volumes at b = 500 s/mm^2^, 500 volumes at b = 1000 s/mm^2^, 2000 s/mm^2^ and 3000 s/mm^2^ each, 600 volumes at b = 4000 s/mm^2^. The data was acquired with a resolution of 2.5 mm isotropic. The MASSIVE dataset was corrected for signal drift (Vos et al., 2017), motion, Eddy current and echo-planar imaging distortions (Leemans & Jones, 2009).

Additionally, we applied our method to the preprocessed data of ten subjects from the Human Connectome Project (HCP). The diffusion MRI data had a resolution of 1.25 mm isotropic and contained 18 volumes at b = 0 s/mm^2^, 90 volumes at b = 1000 s/mm^2^, 2000 s/mm^2^ and 3000 s/mm^2^ each.

An in-house implementation of multi-shell constrained spherical deconvolution (Jeurissen et al., 2014) was used for the FOD estimation.

In order to reconstruct the CST, the seed region was placed close to the internal capsule with 100 points per voxel evenly distributed at a single slice level. The number of seeds per voxel was selected empirically. The motor cortex was segmented as a combination of the precentral and paracentral gyri (Fig. 4) with *FreeSurfer* (Desikan et al., 2006; Fischl et al., 2004; Fischl, 2012) and used as a target region.

**Figure 4:**
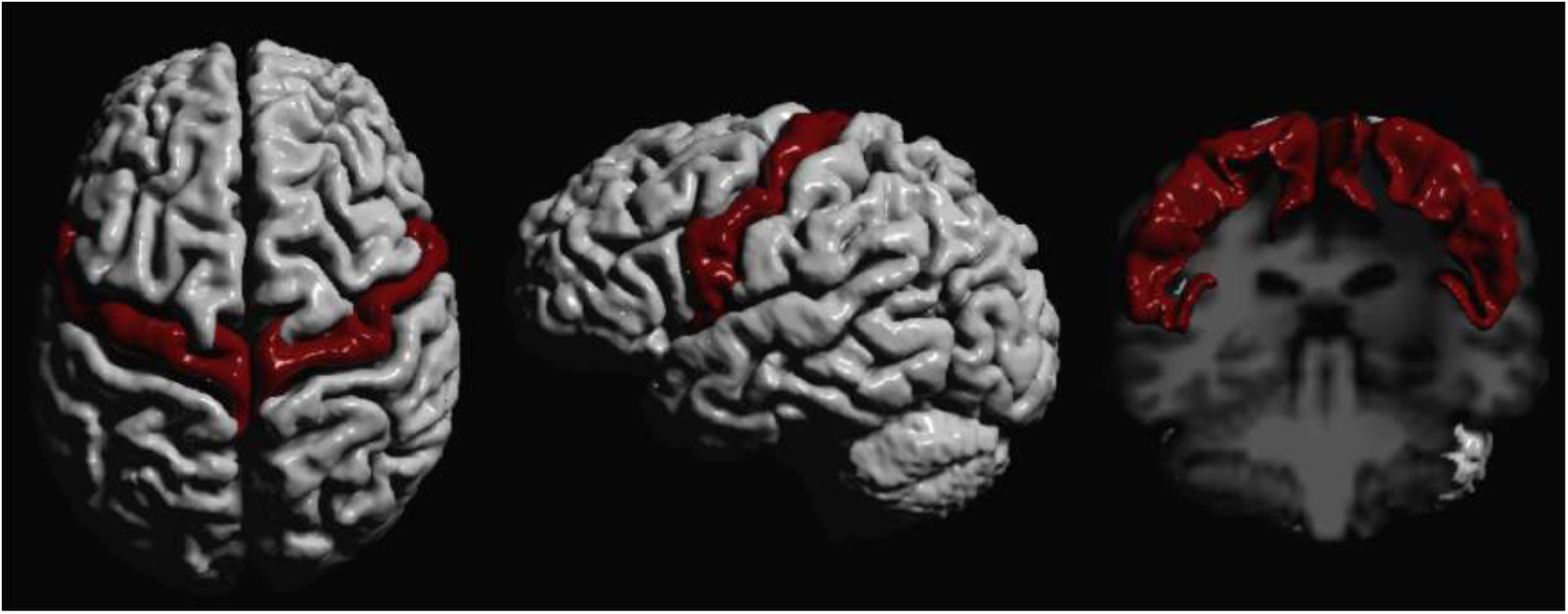
The target cortical region. To reconstruct the corticospinal pathways, the motor cortex (red) was delineated for both hemispheres with *FreeSurfer*.

### 2.3 Experiments

#### Experiment 1. Tractography in silico

As we aim to reach coverage comparable to that achieved by probabilistic approaches while keeping the number of false-positive reconstructions under control, our results are compared against those produced by a probabilistic algorithm, iFOD2 (Tournier et al., 2010), using the noiseless phantom. In all of the experiments the implementation of iFOD2 from the MRtrix package (Tournier et al., 2019) was used and all the options were set to default except for providing the seeding region. One identical seed point was used for tracking in both cases.

The tractography parameters were set as follows: angle threshold = 45^*°*^, maximum order of spherical harmonics *L*_*max*_ = 8, FOD peak value threshold = 0.1, The step size was set to half the voxel size and the number of iterations was set to two. The endpoint regions pictured in Fig. 3 served as target regions of interest for MLFT. They were also used to select the target fibers from the results of iFOD2, which was run with default parameters.

#### Experiment 2. Robustness to noise

The sensitivity of the MLFT to noise was tested. Fiber tracking was performed for the phantoms at varying SNR levels with the same settings as in Experiment 1. The target fibers were then compared across SNR levels.

#### Experiment 3. Tractography in vivo

Our approach was used to delineate the CST with the MASSIVE and HCP brain data described above. Reconstructions with two iterations (for both MASSIVE and HCP) and three iterations (only for MASSIVE) were performed. The motor cortex area of both hemispheres was used as a target region. Other tractography parameter settings were the same as in the previous experiments. The added value of our multi-level strategy was investigated more closely on the example of a fan of the left CST. In the case of the MASSIVE data, the seed regions were placed close to the internal capsule, while for the HCP dataset an axial cross section of the upper part of the brainstem was used for seeding.

In order to see whether MLFT reconstructs parts of the pathways belonging to the corpus callosum (CC), the bundle was delineated with both deterministic CSD-based whole-brain tractography and MLFT. The whole-brain tractography was performed with the same step size and angular deviation threshold as MLFT. The overlap of the CST and the CC generated by MLFT and CSD-based whole-brain tractography, respectively, was visually evaluated.

The results of our algorithm were compared to the results produced by iFOD2 and global tractography (GT) (Christiaens et al., 2015) implemented in MRtrix. Both MLFT and iFOD2 used the same seed regions. Note, iFOD2 was only provided with a mask of a seed region, while MLFT was provided with seeding points as explained above. The streamlines reconstructed by iFOD2 were post-processed to include only the tracts that visit the target cortical area. In the case of GT, the masks of the seed and target regions were used to delineate the CST from the whole-brain tractography. To improve the visual interpretation of the results, implausible streamlines were removed using identical exclusion regions for all methods in case of MASSIVE dataset.

During the analysis of the HCP data, the number of selected pathways was empirically set to 10000 when performing iFOD2 on the HCP data. A NOT gate was used to remove inter-hemispheric connections, due to using common seed region in the brain stem for both of the CST branches.

In order to obtain the whole brain reconstruction with GT, the number of iterations was set to 10^9^, segment length = 1.5mm, maximum spherical harmonics order *L*_*max*_ = 8. The default values were used for the remaining settings.

#### Experiment 4. Topographic organization

Previous research has established that both the motor cortex and the internal capsule can be divided into regions corresponding to specific motor functions, and that such organization is preserved within the CST. (Patestas & Gartner, 2006).The topography preservation index (TPI) (Aydogan & Shi, 2017) was calculated, which highlights whether pathways that pass in close proximity through the internal capsule also have closely located endpoints in the motor area. Thus, the index reflects how well the internal organization is preserved in the bundle reconstruction. To visually appreciate such organization, the CST streamlines were colored according to the part of the area of the motor cortex they reach. This allows to visually check whether the pathways reconstructed by MLFT and iFOD2 on the MASSIVE data corresponded to the anatomical position of the same function in the internal capsule.

#### Experiment 5. Anatomical plausibility

Pathways’ coherence was evaluated to determine the geometric similarity between pathways closely located to each other along their length. To this end, the minimum average direct-flip (MADF) distance was employed, which previously has been used in bundle clustering applications (Visser et al., 2011; Garyfallidis et al., 2012). This metric represents the average point-to-point distance between two pathways. It is defined in the following way: 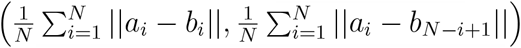, where *a*_*i*_ and *b*_*i*_ are the points of the pathways *A* and *B* of length *N*, respectively. This metric requires the compared tracts to contain an equal number of points, which is why all the pathways were uniformly resampled to *N* = 200 points. Evaluations were performed on the left and right CST bundles of the MASSIVE and HCP data obtained by the tested methods without filtering gates. For each set of reconstructed pathways of a given subject, an all-to-all distance matrix was calculated. Then, for each pathway, the minimum distance was calculated based on that matrix.

## 3 Results

### Experiment 1. Tractography in silico

Both MLFT and iFOD2 reconstructed all the phantom branches of the noiseless DWI phantom, as shown in Fig. 5. It can be observed that the results of MLFT follow the underlying simulated directions, whereas iFOD2 produces trajectories oscillating around the ground truth.

**Figure 5:**
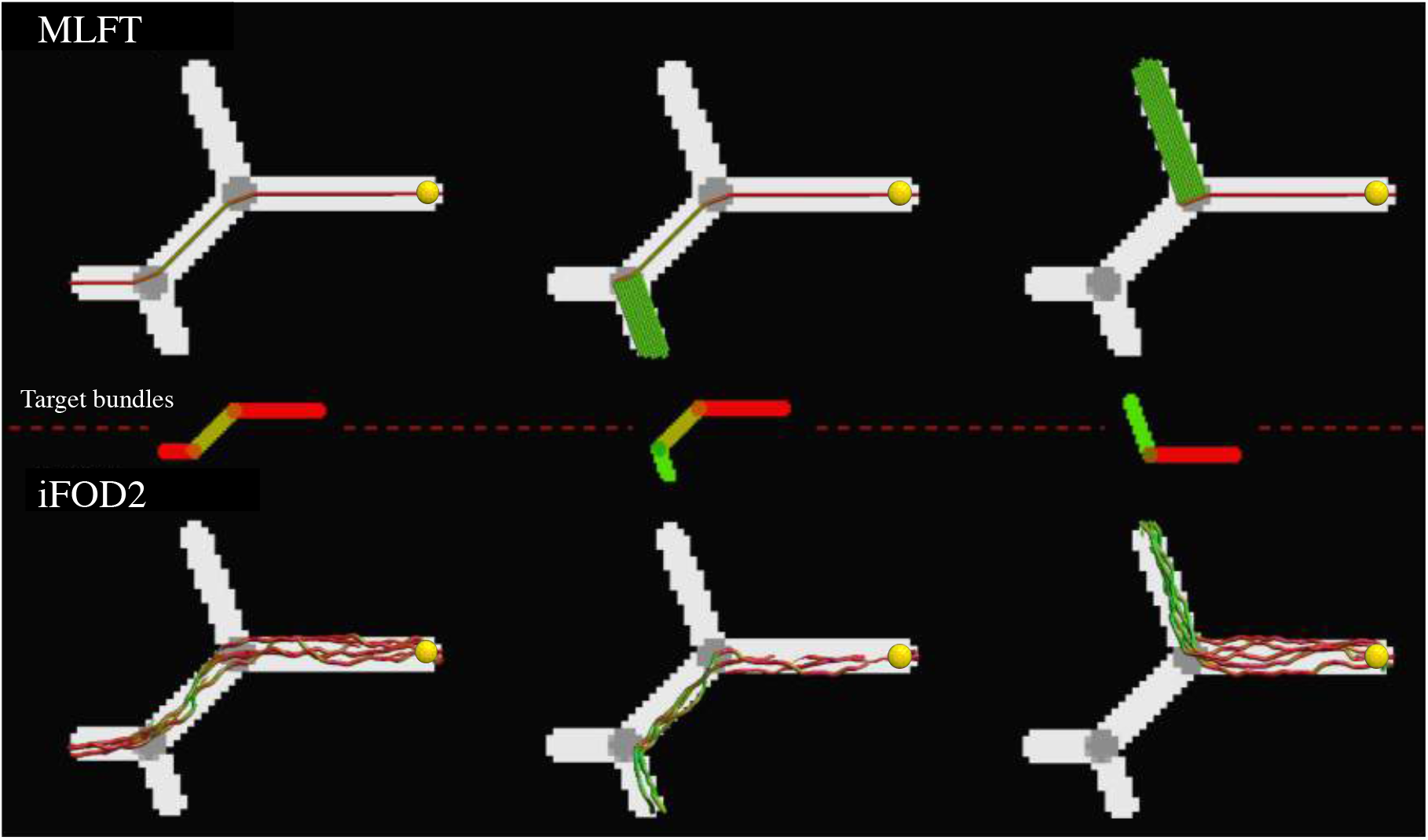
Performance of the considered methods in phantoms (FA map). The top row shows the results of MLFT and the bottom row those of the iFOD2 algorithm. The middle row illustrates the target fibers per column (colored FA map). The same single seed point (yellow sphere) was used for both algorithms. The results of iFOD2 were subsampled for easier visual assessment. Streamlines’ colors as well as colored FA map are based on orientation color-coding.

### Experiment 2. Robustness to noise

The results of MLFT obtained for three different SNRs are presented in Fig. 6. A slight misalignment lower than 10^*°*^ can be observed at the branching point at SNR = 25, which becomes more evident at SNR = 15 with values up to 30^*°*^. The case with the lowest SNR is also characterized by an increased pathway splitting at the points where original bundles diverge, as can be seen in the top row in Fig. 6.

**Figure 6:**
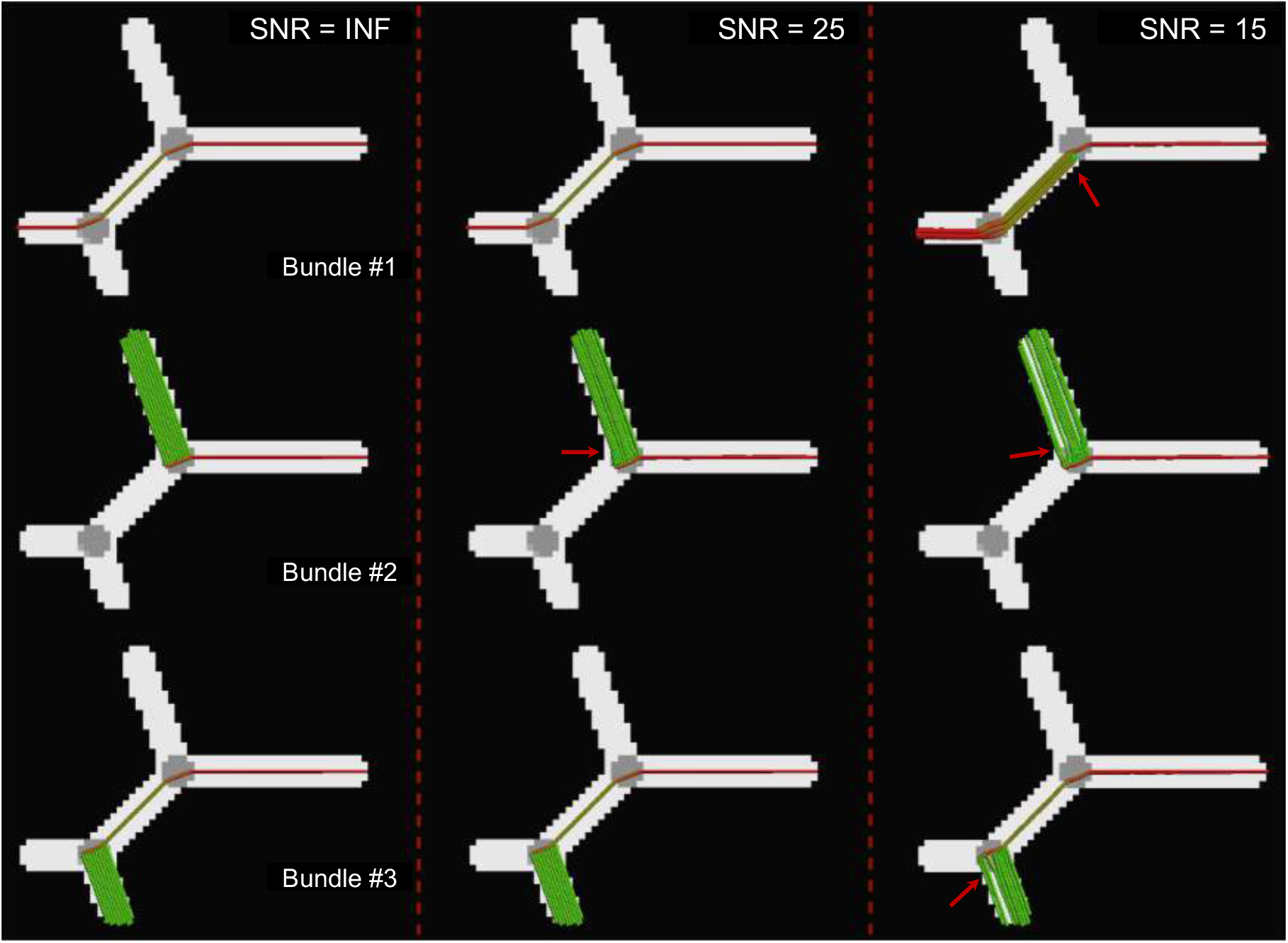
Tracts reconstructed by MLFT on the phantom data (FA map) at multiple SNR levels. Considerable angular errors are only observed at SNR = 15: increased splitting and direction perturbations up to 30^*°*^(red arrows). At SNR = 25 there is a minor angular deviation below 10^*°*^(red arrow). Streamlines are colored using standard orientation color-coding.

### Experiment 3. Tractography in vivo

The multi-level structure of the reconstructed left CST bundle can be seen in Fig. 7, which clearly shows the benefit of the proposed algorithm over conventional deterministic CSD-based tractography with the improved extent of the bundle fanning. The addition of an extra layer, shown in Fig. 8, increases the number of streamlines reaching the motor cortex but does not bring further improvement to the coverage of the motor cortex. Consequently, in all of the in-vivo experiments the number of levels was set to two.

**Figure 7:**
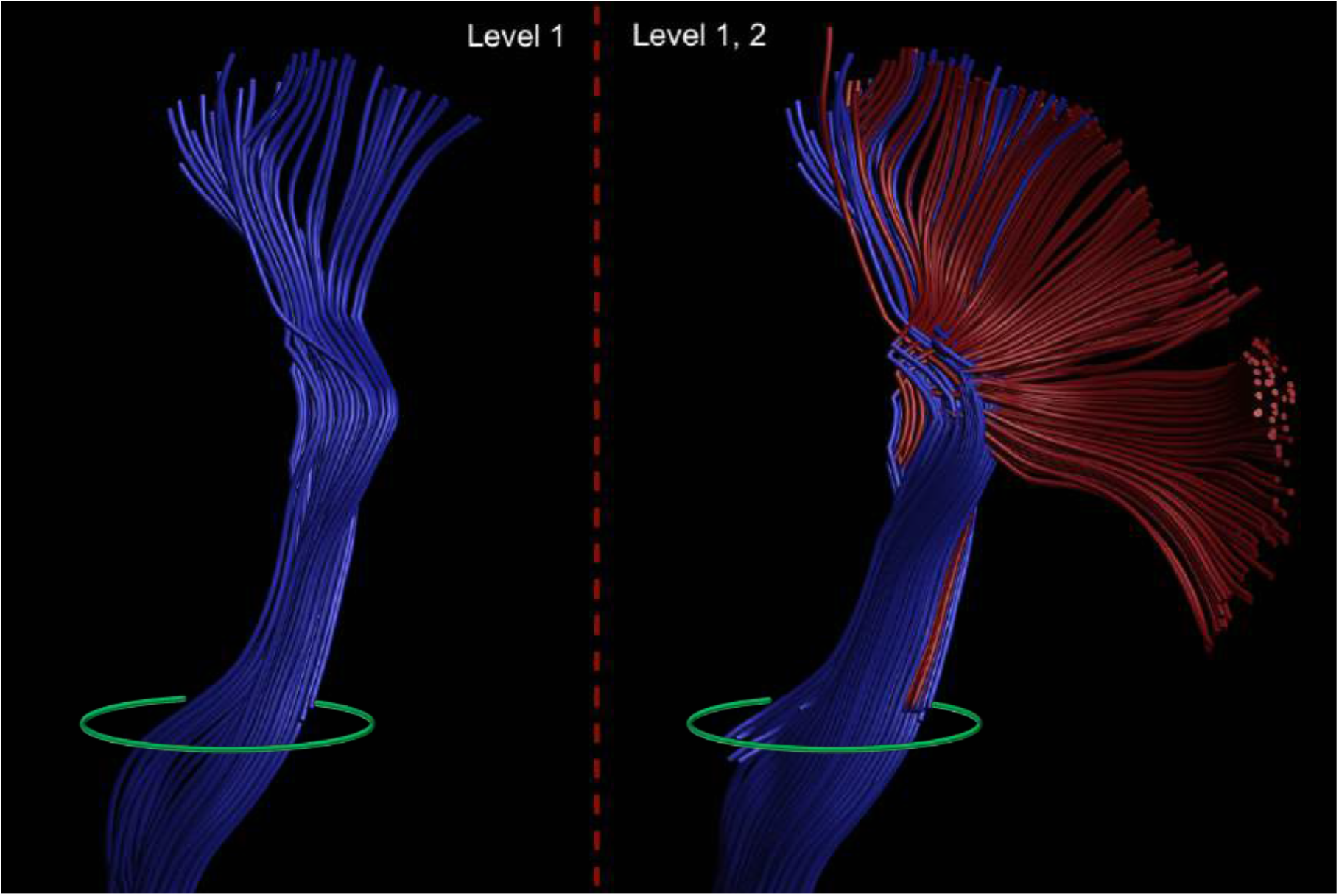
Fiber pathways reconstructed by the deterministic CSD-based approach (left) and MLFT (right) from the same seed region (green). Adding the second-level branches (red) to the pathways obtained at the first level (blue) improves the extent of the reconstructed bundle.

**Figure 8:**
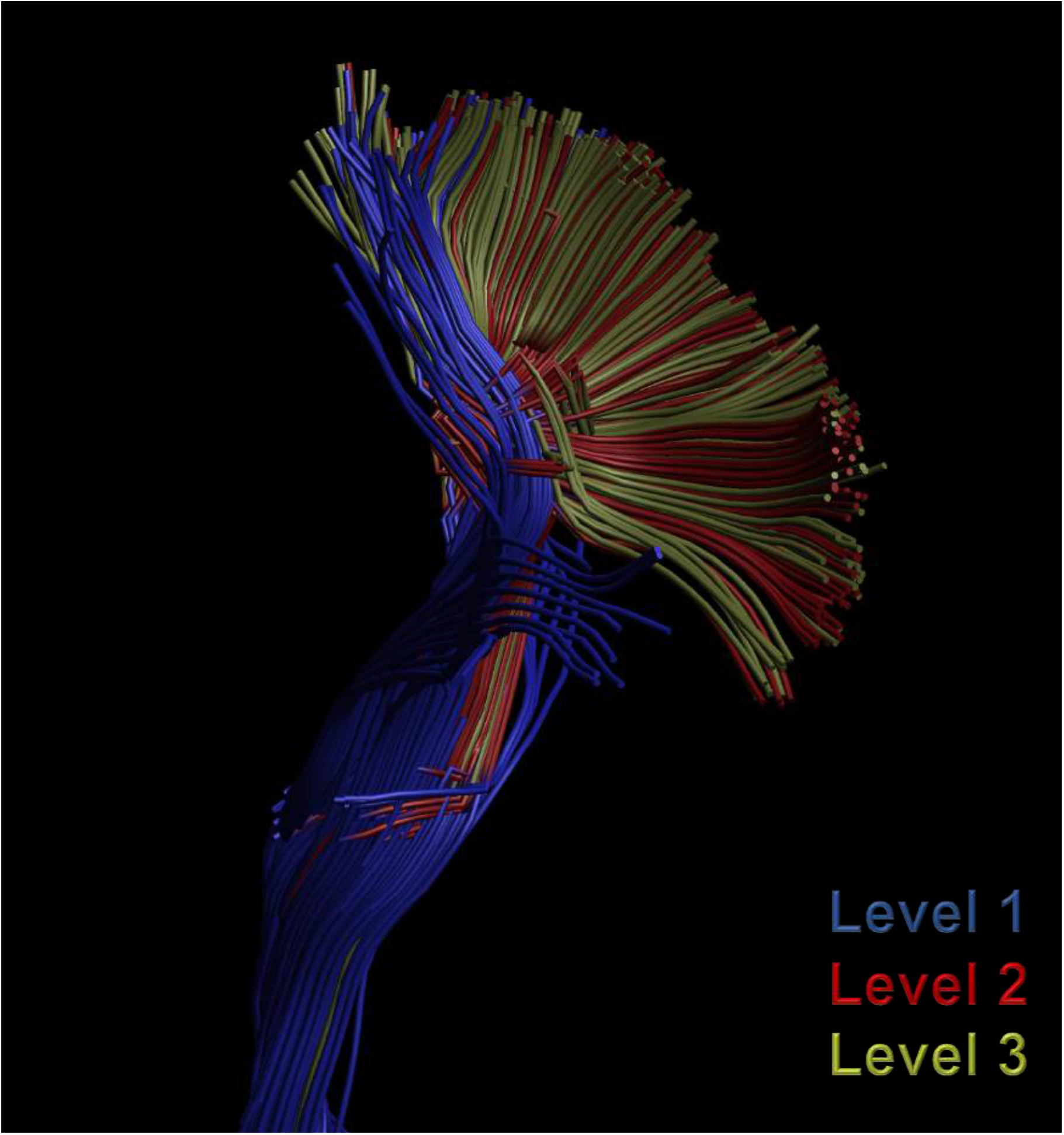
The 3-level reconstruction of the left CST by MLFT using the MASSIVE data from the same seed region shown in Fig. 7. The result does not show coverage improvements over the 2-level reconstruction.

The full reconstruction of the CST segmented by MLFT in the MASSIVE data is shown in Fig. 9. It can be observed that the obtained pathways densely cover most of the motor cortex. Further, neighboring streamlines are geometrically similar. This contrasts with the result produced by the probabilistic approach pictured in Fig. 10, which shows pathways with high tortuosity as well as multiple spurious fibers. At the same time, both MLFT and iFOD2 cover most of the motor area. In the iFOD2 reconstruction, the pathways traversing into contralateral hemisphere are present due to using visitation-based inclusion criteria at the post-processing stage, which does not filter out pathways bending after visiting the target region and propagating through the CC.

**Figure 9:**
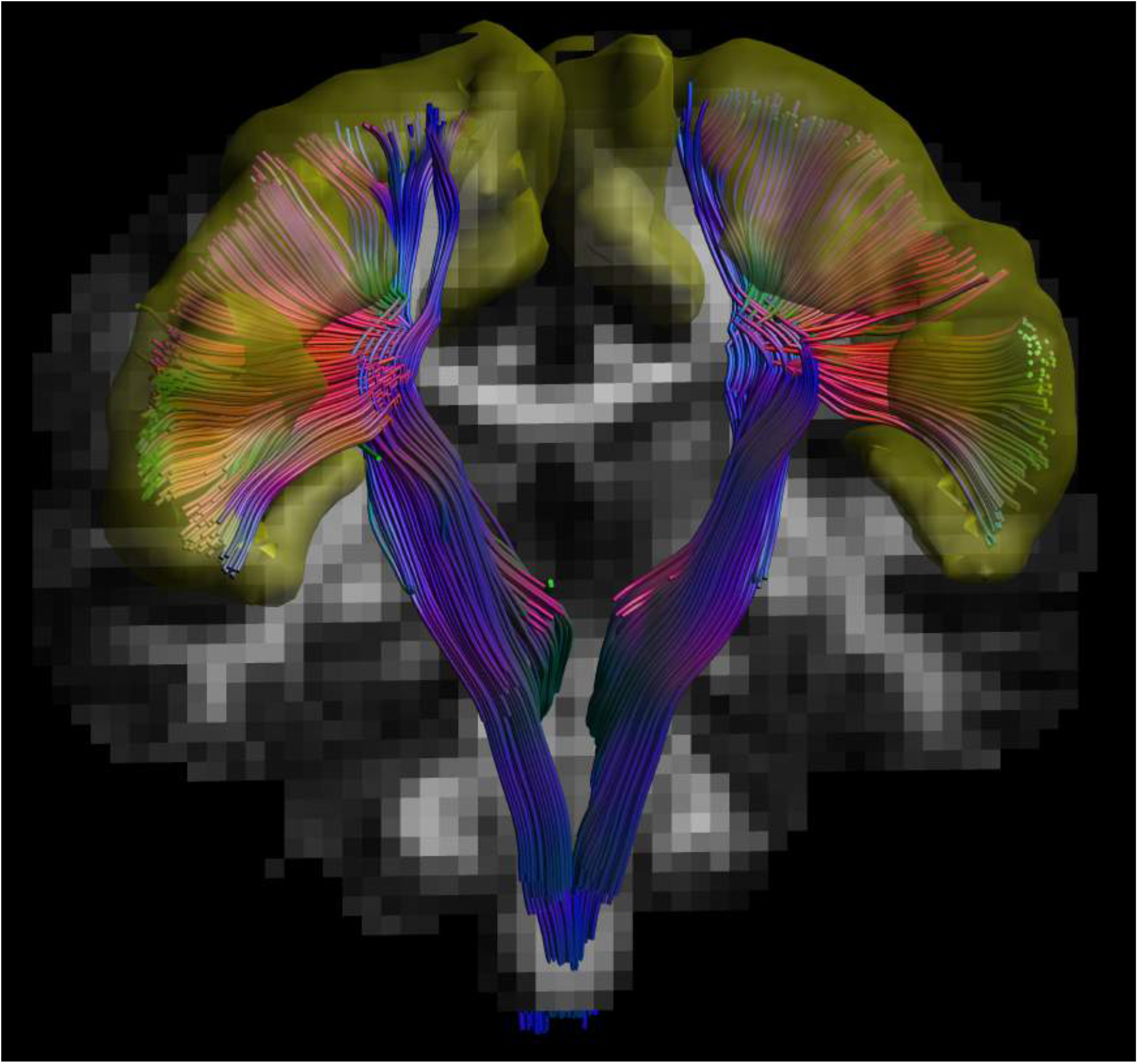
Corticospinal pathways reconstructed by MLFT using the MASSIVE data. The motor cortex is shown in yellow.

**Figure 10:**
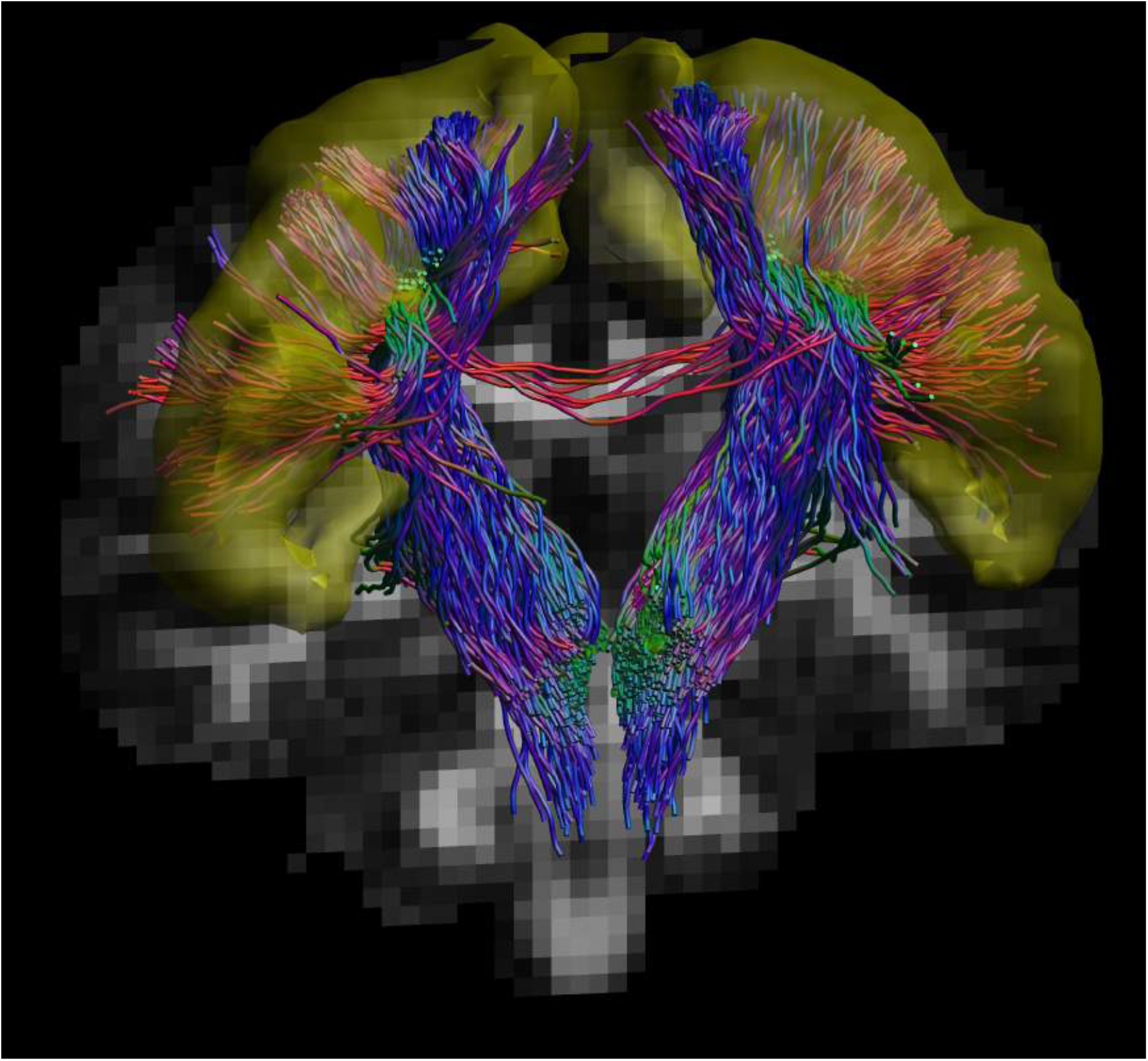
Corticospinal pathways reconstructed by iFOD2 using the MASSIVE data. The motor cortex is shown in yellow. Some of the pathways enter the motor cortex and diverge into the CC propagating into the contralateral hemisphere.

The reconstruction achieved by GT using the MASSIVE data is pictured in Fig. 11. Although the CST fanning is quite sparse, it reaches all the main parts of the motor cortex. The sparsity allows for a closer comparison of the multi-level and global tractography results which can be seen in Fig. 12. Unlike GT, the CST reconstructed by MLFT does not reach the approximate leg-related motor area. In the face area the pathways generated by GT are aligned to those generated by MLFT, although they do not show any branching, but rather a smooth curving trajectory.

**Figure 11:**
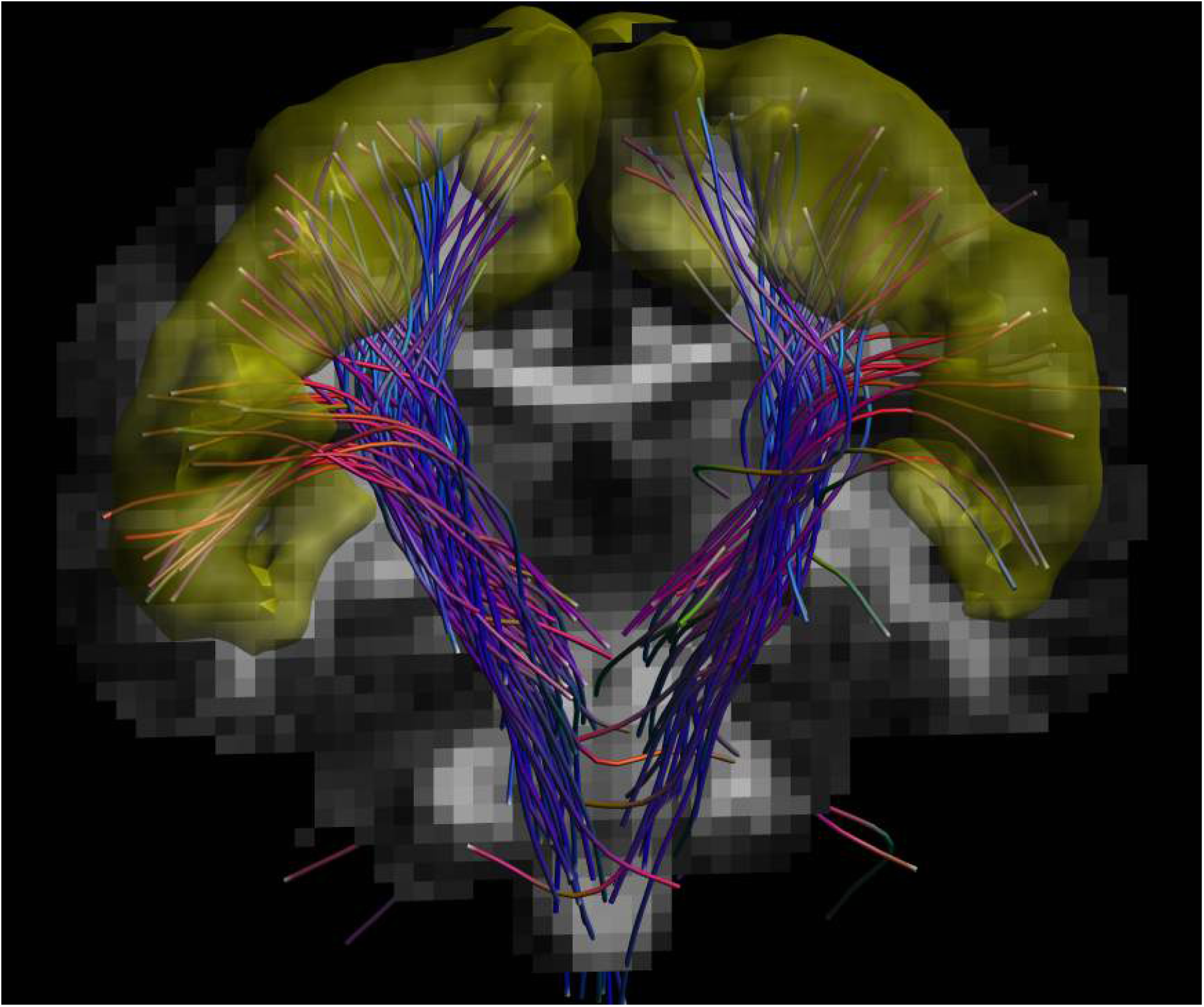
Corticospinal pathways reconstructed by GT using the MASSIVE data. The motor cortex is shown in yellow.

**Figure 12:**
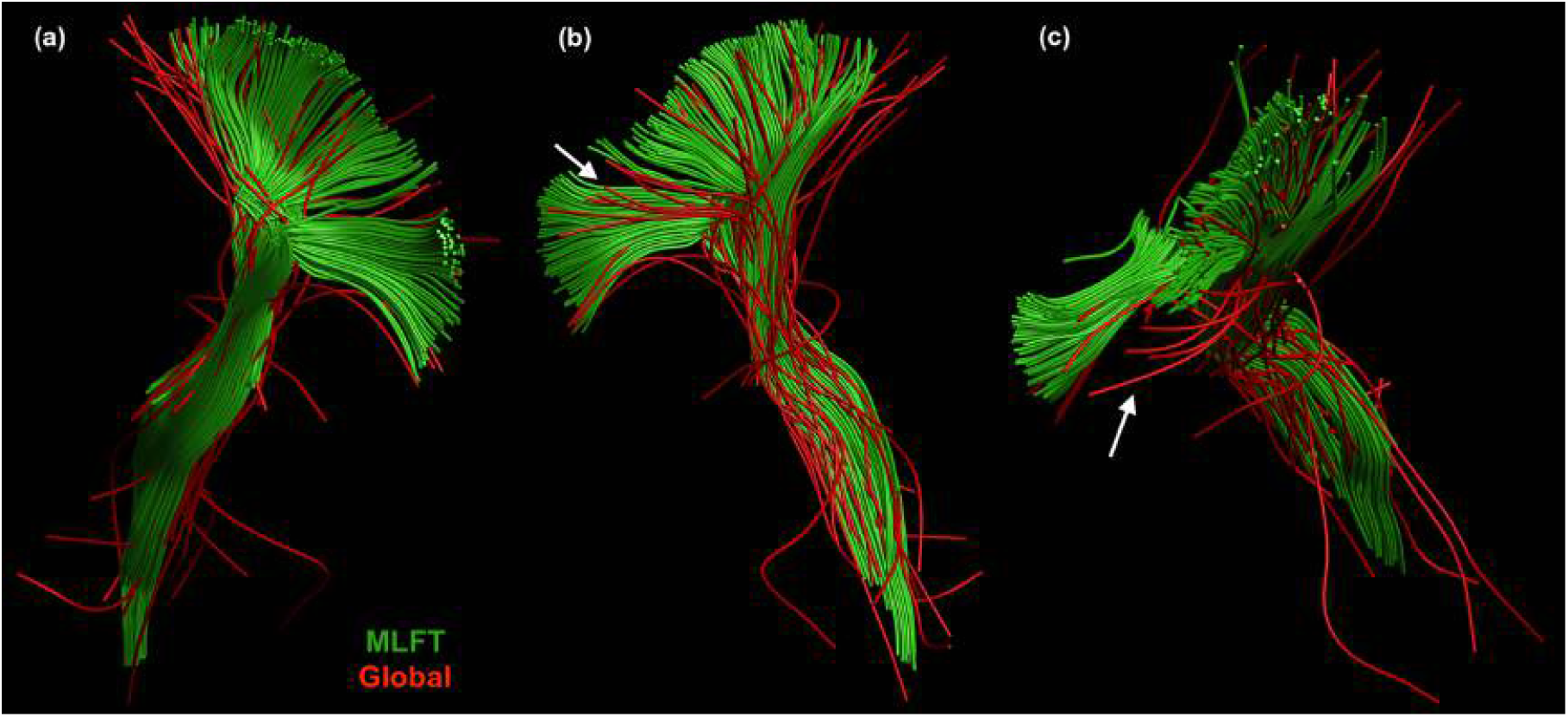
Comparison of the left CST reconstructions obtained by MLFT (green) and GT (red). (a) The reconstruction by GT is sparser, but it provides additional coverage towards the approximate leg area unlike the MLFT reconstruction. (b) Pathways delineated by GT generally follow the same trajectory of the bundle reconstructed by MLFT but with smoother branching turns. (c) Some of the GT-produced pathways that are reaching the face motor area (white arrows in (b) and (c)) are shifted towards posterior part of the brain, and are not completely aligned with others as well as with the corresponding part of the MLFT-reconstructed fanning.

The CST bundles that were reconstructed for the HCP subjects by the proposed approach, iFOD2 and GT are shown in Fig. 13, Fig. 14 and Fig. 15, respectively. Regarding iFOD2, the results have the same characteristics as the results obtained using the MASSIVE data described above. Generally, both MLFT and iFOD2 reconstructions are represented by the bundles with a plausible fanning extent. In addition, the bundles produced by MLFT are more well-aligned to each other than the results of iFOD2. At the same time, GT underperformed in terms of the extent compared to its result using the MASSIVE data.

**Figure 13:**
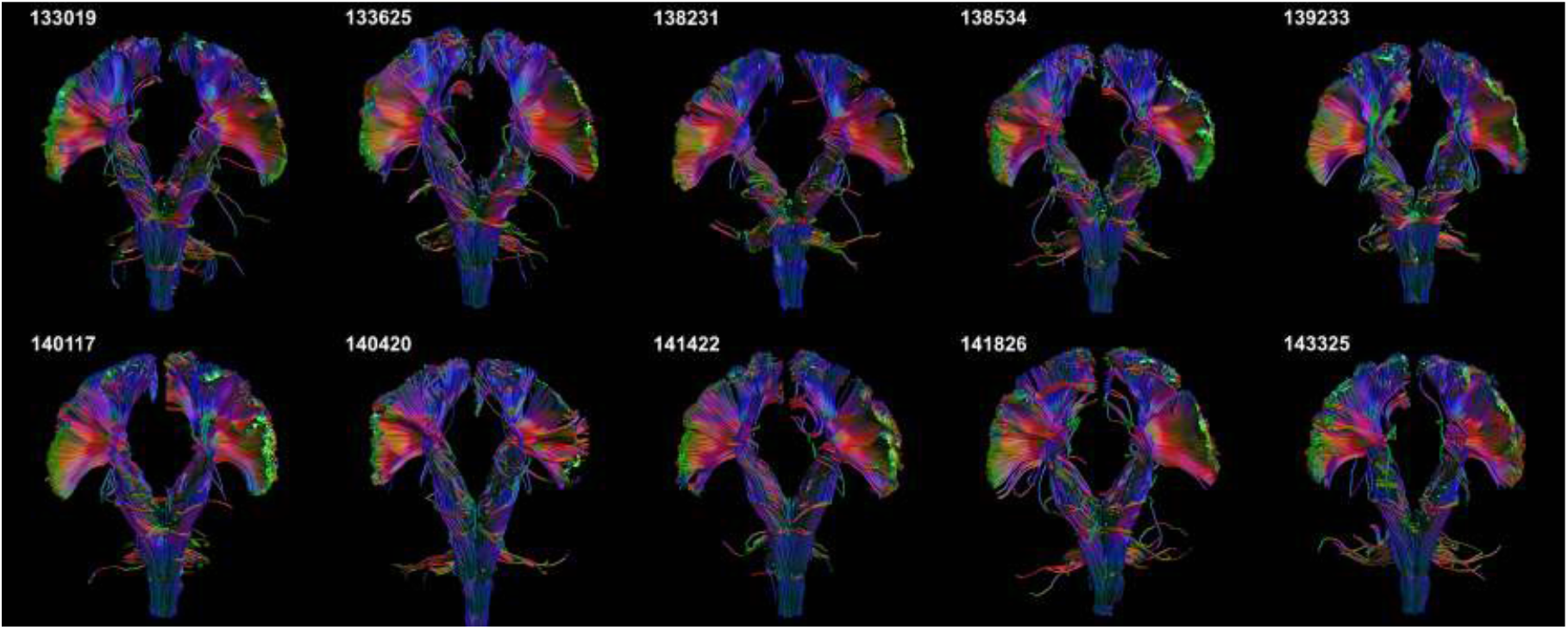
The CST reconstructions obtained by MLFT using the HCP data. The reconstructed bundles are inline with the observations in Fig. 9 and consistent with each other.

**Figure 14:**
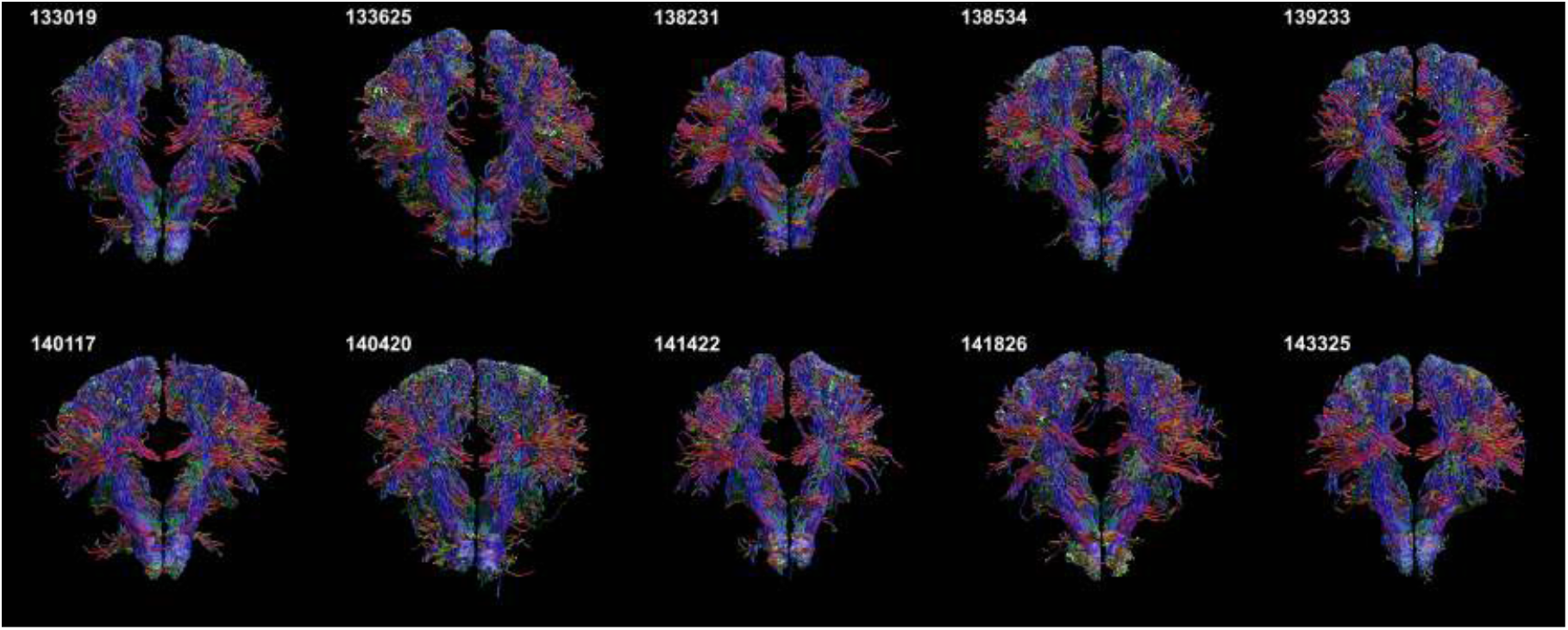
The CST reconstructions obtained by iFOD2 using the HCP data. The radial extent is comparable to the results obtained by MLFT (Fig. 13).

**Figure 15:**
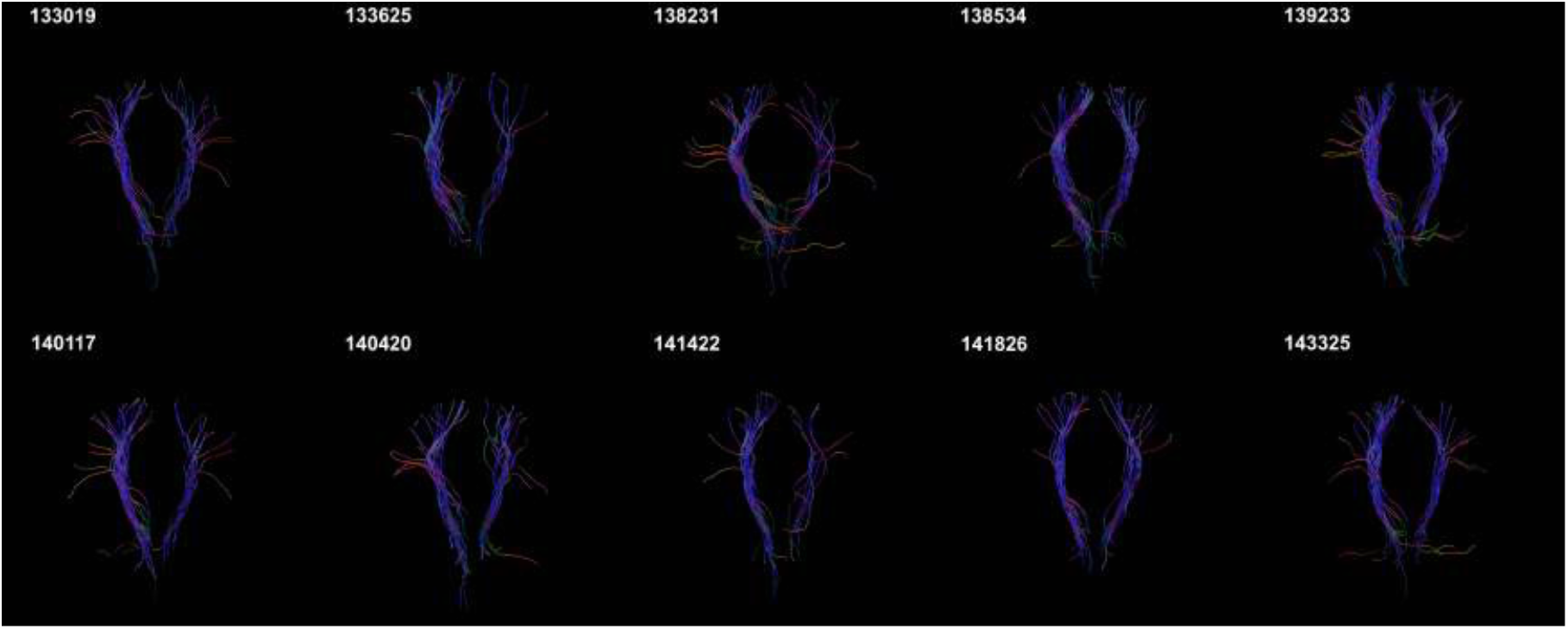
The CST reconstructions obtained by GT using the HCP data. The radial extent of some of these reconstructions is comparable to the ones obtained by MLFT (Fig. 13) but with less satisfactorily spatial coverage.

The CST and CC bundles reconstructed using the MASSIVE data are depicted in Fig. 16 for comparison. In the sagittal plane it is well visible that most of the CST fanning does not overlap with the CC pathways, as the CC is not covering most of the motor cortex.

**Figure 16:**
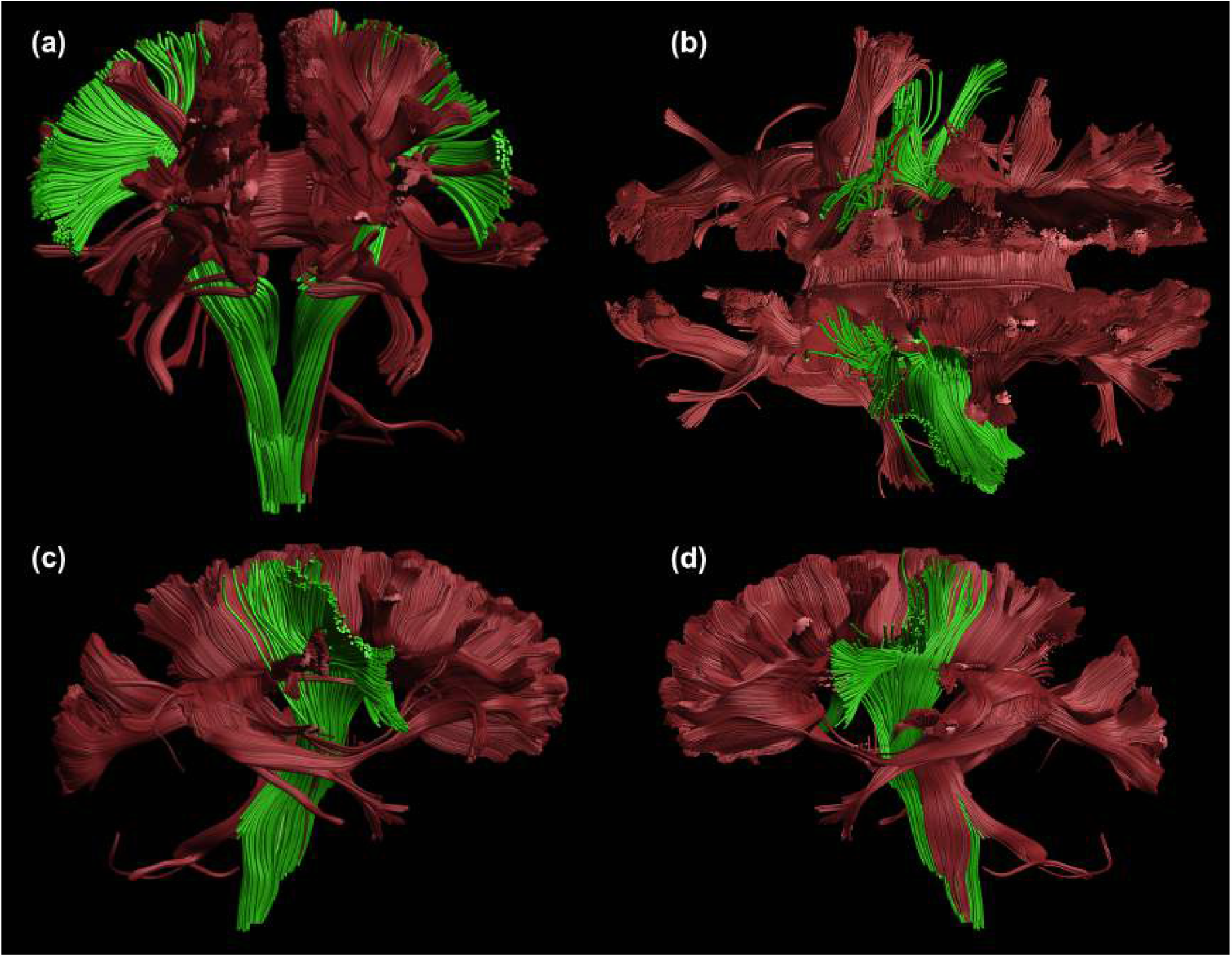
The CST bundle (green) reconstructed by MLFT and the CC bundle (red) reconstructed by whole-brain CSD-based tractography using the MASSIVE dataset. The coronal (a) and axial (b) projections show little overlap of the bundles supporting the fact that MLFT does not reconstruct crossing bundles as branches. This is also confirmed from both right (c) and left (d) sagittal projections.

The original reconstruction of the whole CST by MLFT using the MASSIVE data without using exclusion gates can be found in supplementary materials (Fig. S1).

Although this work focuses on the CST reconstruction we have additionally evaluated the generalizability of MLFT using the MASSIVE dataset by reconstructing the cingulum bundle. The results can be found in the supplementary materials (Fig. S2).

### Experiment 4. Topographic organization

The TPI scores are reported in Table 1. MLFT outperforms other algorithms showing lower values of the TPI metric, and thus better coherence, for every subject. In this regards, iFOD2 and GT show generally comparable performance to each other.

**Table 1:** Topography preservation indices of the left and right CST reconstructions by MLFT, iFOD2 and GT (the lowest score is indicated in bold). The MLFT-related indices are consistently the lowest for both CST branches.

| Subject | Left |  |  | Right |  |  |
| --- | --- | --- | --- | --- | --- | --- |
|  | MLFT | iFOD2 | Global | MLFT | iFOD2 | Global |
| MASSIVE | **0.08** | 0.15 | 0.18 | **0.08** | 0.21 | 0.17 |
| 133019 | **0.06** | 0.13 | 0.36 | **0.05** | 0.1 | 0.21 |
| 133625 | **0.05** | 0.36 | 0.11 | **0.05** | 0.1 | 0.08 |
| 138231 | **0.06** | 0.36 | 0.2 | **0.04** | 0.09 | 0.25 |
| 138534 | **0.05** | 0.3 | 0.10 | **0.05** | 0.16 | 0.23 |
| 139233 | **0.07** | 0.43 | 0.12 | **0.07** | 0.15 | 0.1 |
| 140117 | **0.05** | 0.11 | 0.13 | **0.04** | 0.1 | 0.11 |
| 140420 | **0.07** | 0.15 | 0.14 | **0.06** | 0.1 | 0.15 |
| 141422 | **0.06** | 0.46 | 0.09 | **0.04** | 0.09 | 0.26 |
| 141826 | **0.05** | 0.18 | 0.12 | **0.05** | 0.09 | 0.1 |
| 143325 | **0.04** | 0.08 | 0.13 | **0.04** | 0.1 | 0.09 |

Fig. 17 shows the pathways color-coded according to their final locations in the motor cortex. The visualization demonstrates that MLFT maintains the anatomical configuration of the pathways, according to which the organization of tracts connecting specific sub-domains of the motor cortex is maintained throughout the bundle (Patestas & Gartner, 2006). In contrast, the bundle produced by iFOD2 is less organized.

**Figure 17:**
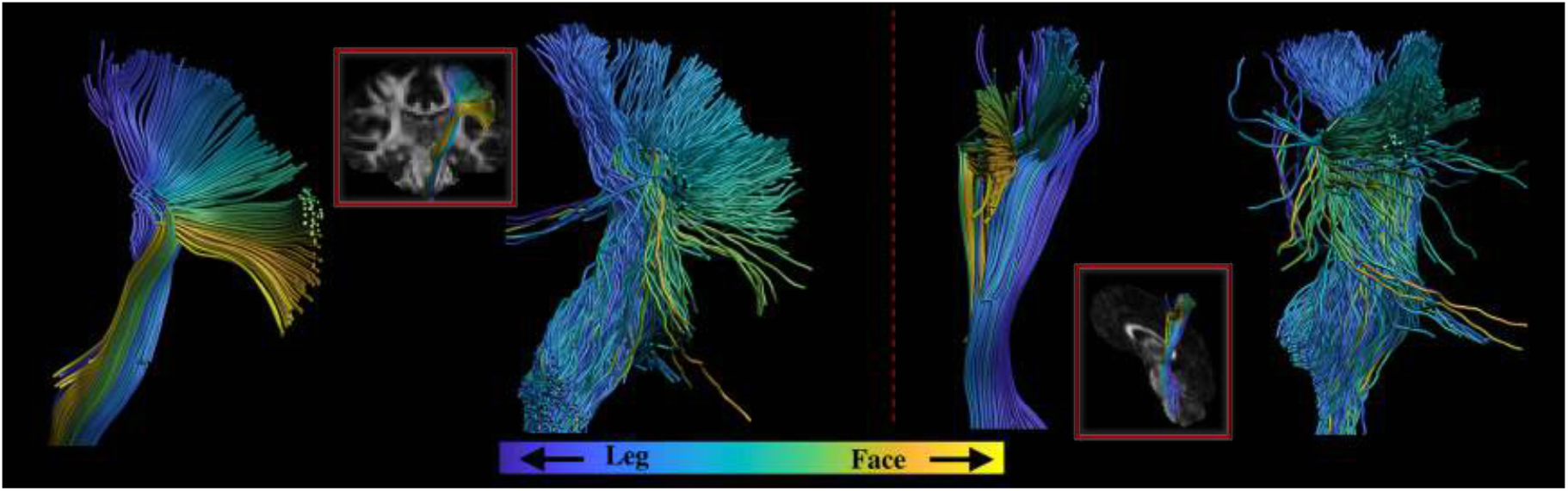
Coronal and sagittal views of the left CST reconstructed by MLFT and iFOD2 using the MASSIVE data. The fiber pathways are colored according to the locations of their endpoints in the motor cortex. The pathways reconstructed by MLFT are shown to have a clearer topographic organization.

The color-coded topographic organization of the MLFT result without filtering can be found in the supplementary materials (Fig. S3).

### Experiment 5. Anatomical plausibility

The normalized histograms of the MADF distance are shown in Fig. 18. The distributions are similar across subjects per tractography approach and show that the distance between the closest pathways obtained by MLFT is generally smaller than that of iFOD2. The results of GT showed the highest distance, which is attributed to the sparsity of the bundles.

**Figure 18:**
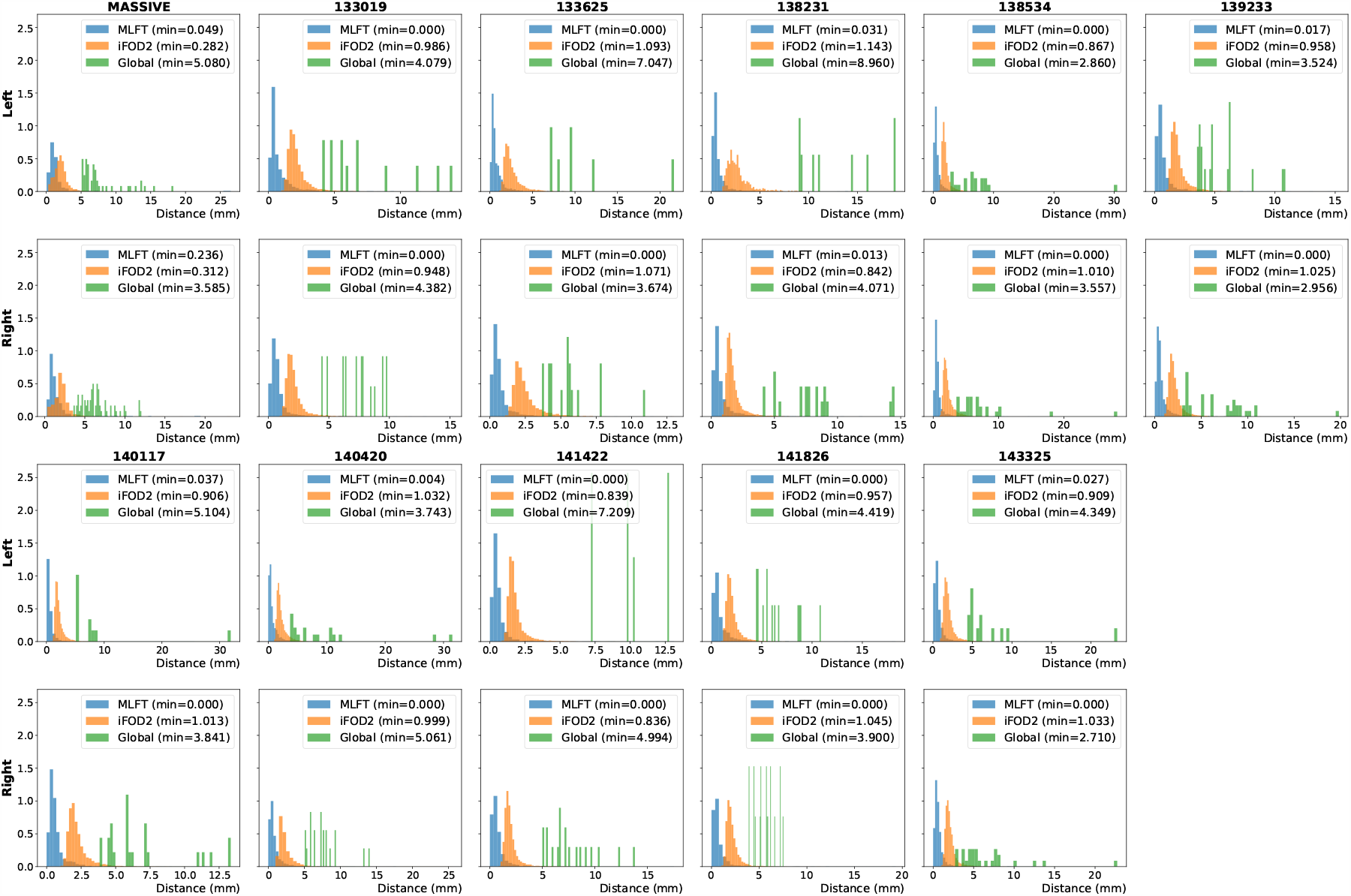
Normalized distributions of the distances from each pathway to the nearest neighbor based on the MADF metric for all of the processed subjects. The distributions of the distances appear to be similar across subjects.

## 4 Discussion

In this study we introduced a novel strategy to perform fiber tractography that combines the best features of probabilistic and deterministic tracking algorithms. The strategy we propose achieves anatomically plausible reconstructions of the CST bundles, is robust and reproducible, and maintains topographic organization. Each iteration of the proposed tractography algorithm adds extra branches, which might reveal pathways that were neglected in the previous step. By allowing for pathway branching, our method might address the fact that axon branching can be observed in the brain (O’Leary & Terashima, 1988). Given that image resolution is usually not sufficient to distinguish branching points, some of the FOD peaks might not only be an indication of crossing fibers, but of splitting ones. For that reason all the peaks should be processed to reconstruct a branch of the same bundle that would have been missed otherwise.

Given the improved extent of the bundles and anatomically imposed control over false positives, our approach is attractive for a number of applications. It can be used for presurgical planning as it reveals more pathways than DT-based tractography (Farquharson et al., 2013; Fortin et al., 2012), which is often used in clinical practice, without overwhelming the user with anatomically inaccurate pathways. Robust tractography is also relevant in case of chronic stroke (Jones, 2010; Pierpaoli et al., 2001), which requires an indication of abnormality occurrence in the CST bundle. The same holds for the study of multiple sclerosis (Jones, 2010; Pagani et al., 2005; Reich et al., 2007) and epilepsy (Jones, 2010; Kreilkamp et al., 2017).

### 4.1 MLFT features

With simulations, we have shown that MLFT reconstructs branching fiber configurations that are less tortuous and erratic as compared to the probabilistic algorithm (Fig. 5). Additionally, the results of MLFT are completely reproducible because new step directions are chosen in a deterministic manner during pathway propagation.

Robustness to noise is another important aspect to consider when comparing to probabilistic tractography, as this is reportedly highly sensitive to SNR level (Mesri et al., 2016). In order to analyze the sensitivity of our algorithm, the same phantom bundles were simulated with three different SNR levels. The changes in the reconstructed pathways became clearly visible only at the lowest SNR level (SNR = 15), being reflected in an increased splitting at branching points and occasional perturbations after branching (Fig. 6).

We have shown that for in-vivo images the concept of selecting peaks produces improved tractography results in a number of aspects. The reconstructed CST bundles for both hemispheres are characterized by smooth, well-aligned pathways, as they would in case of deterministic CSD-based tractography. An additional benefit is that the fanning close to the motor cortex is well delineated (Fig. 7). Thus, despite comparable reconstructions of the CST fanning with both MLFT and iFOD2, the results produced by iFOD2 show an erratic behavior including multiple spurious tracts (Figs. 9, 10, 13 and 14).

Most of the fanning consists of the second level branches, which might often look as if they diverge into another bundle at the branching points making a sharp turn. However, high angular deviations have been observed by Wedeen et al. (2012). Similarly, Mortazavi et al. (2018) also observed axon T-branching as well as sharp turns at sub-millimeter scale performing tract tracing experiments. Both of those papers are based on the analysis of the macaque brain, but the statements are also valid for the human brain, which is reportedly congruent to the structure of the macaque brain (Mortazavi et al., 2018).

Discussing the validity of the reconstructions our method provides, we can also refer to Fig. 16 at this point. It is known that part of the CC originates from the motor cortex (Wassermann et al., 2016; Witelson, 1989). Thus, a successful reconstruction of the motor part of the CC remains prone to ambiguity as the CST pathways are present in that area as well. Advocating for the validity of our reconstruction, the shapes of the MLFT-recontructed bundles are also similar to those presented by Wasserthal et al. (2018). Additionally, the comparison to the results of GT (Fig. 12) has shown that this alternative approach reconstructs similar pathways, although with certain smoothing of the high-angular bifurcations that are observed in MLFT results. In general, the resemblance between the second level of the CST and the CC bundle can be explained by the co-alignment of the pathways of different bundles under the motor region which was reported by Krieg (1955) for the macaque brain and for the human brain. This does not prove that the similar pathways are true positive but serves as a reference which shows stable delineation of certain structures across various algorithms.

MLFT well preserves topographic organization, as can be seen in Fig. 17. This is also reflected in the comparison of the TPI scores (Table 1) across all the subjects analyzed in this work. By following the FOD peaks, we propagate the streamlines into the most certain direction of diffusion and, consequently, are less susceptible to noise effects. This leads to a more stable pathway propagation and, consequently, to a better organization of the bundle, as also supported by the presented high pathway coherence (Fig. 18).

### 4.2 Limitations

Some degree of uncertainty propagates in the results from the CSD procedure, as the response function is not voxel-wise perfect and FOD peaks have limited angular resolution. This limitation is, however, inherent to most tractography algorithms. Further, branching along a pathway might generate false-positive reconstructions or intersect crossing bundles, although our results suggest this is limited, likely due to the incorporation of correctly chosen anatomical priors to control the rate of false-positive pathways.

An accurate seeding area and a sufficient subsampling are two other components that a good reconstruction depends on. Excessively sparse seeding might lead to the absence of the expected branching structures in complex cases. One of the recommendations would be to set the seeding regions using the atlas-based gates (ROIs) for bundle delineation (Catani & Thiebaut de Schotten,2008;Wakana et al.,2007). Another approach would be to use cortical masks as seeds and gates as target regions. However, this might increase the computational load as multiple bundles may be connected to a single cortical region. The first iteration would be then polluted with obviously false-positive pathways that would be used as seeds for the next iteration. Given that in the worst case the proposed algorithm has exponential complexity, heuristics implementing on-the-fly pathway filtering could reduce the processing time and make this approach viable.

It must be noted that in this work we did not focus on devising an approach for estimating the required number of levels based on convergence criteria, which might be useful for clinical translation. For this study, those settings were identified empirically.

### 4.3 Future work

Although in this work we combined our proposed concept with the deterministic tractography algorithm, it is generic enough to be combined with other approaches. For example, it can also be combined with probabilistic approaches by representing the FODs as mixture distributions.

## 5 Conclusion

Multi-level fiber tractography is a new fiber tractography method that reconstructs fiber pathways with better extent than deterministic methods while avoiding the false-positive issues of probabilistic methods. MLFT addresses the reconstruction of branching pathways, which, to date, has been poorly investigated to the best of our knowledge. Thanks to its robustness and the preservation of topographic organization, our approach may assist in clinical practice as well as in virtual dissection studies.

## 6 Acknowledgments

Andrey Zhylka is supported by the European Union’s Horizon 2020 research and innovation program [grant number 765148]. Alberto De Luca is supported by EraNet-NEURON RepImpact project [grant number R4195]. Data were provided in part by the Human Connectome Project, WU-Minn Consortium (Principal Investigators: David Van Essen and Kamil Ugurbil; 1U54MH091657) funded by the 16 NIH Institutes and Centers that support the NIH Blueprint for Neuroscience Research; and by the McDonnell Center for Systems Neuroscience at Washington University. Declarations of interest: none.

---

**Figure S1:**
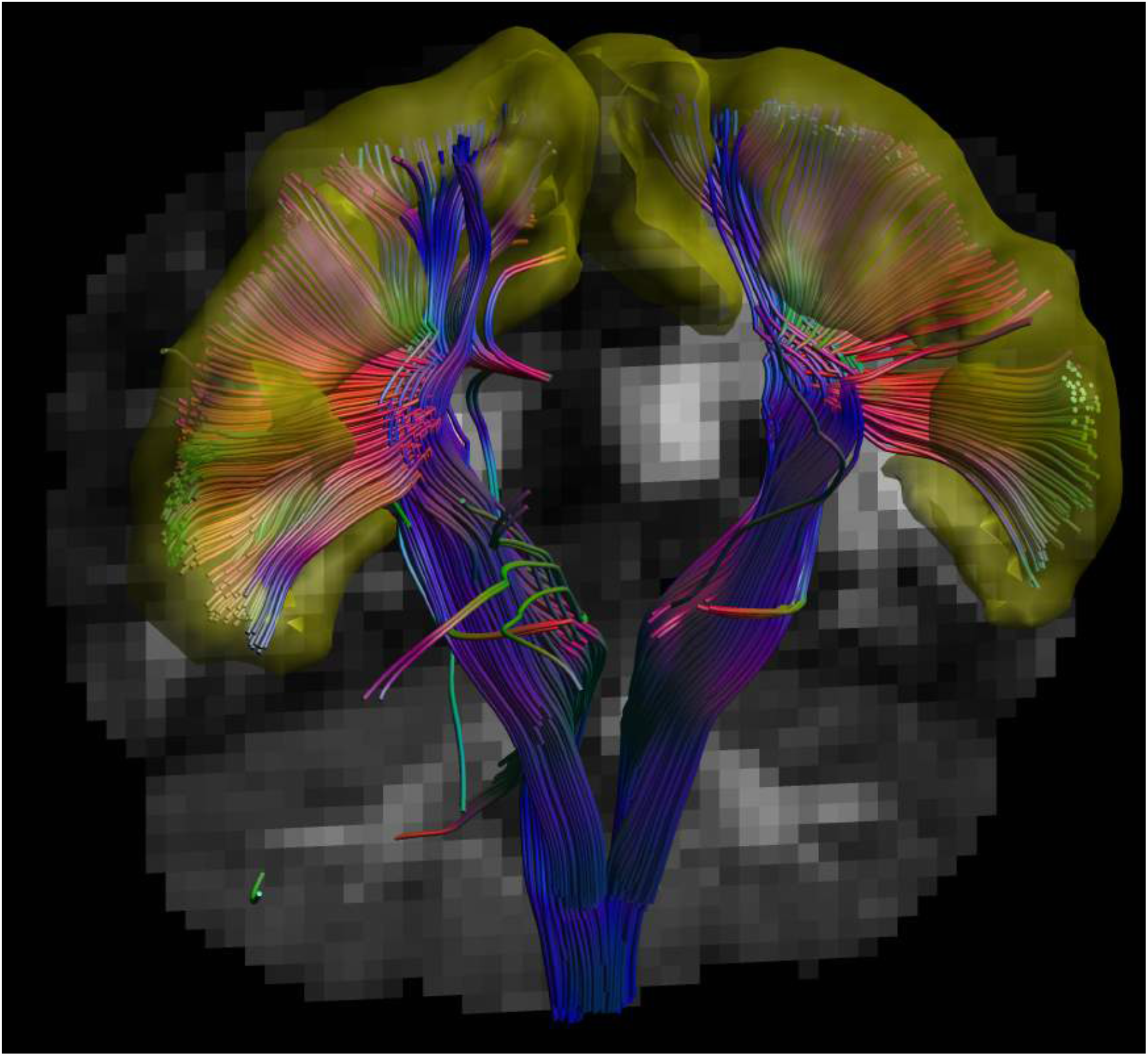
The whole CST reconstructed by MLFT using the MASSIVE data without consecutive filtering with exclusion gates. The motor cortex is rendered in yellow. The residual false positives are mainly due to placing the seed region around internal capsule and not imposing the constraint for pathways to enter the brain stem.

**Figure S2:**
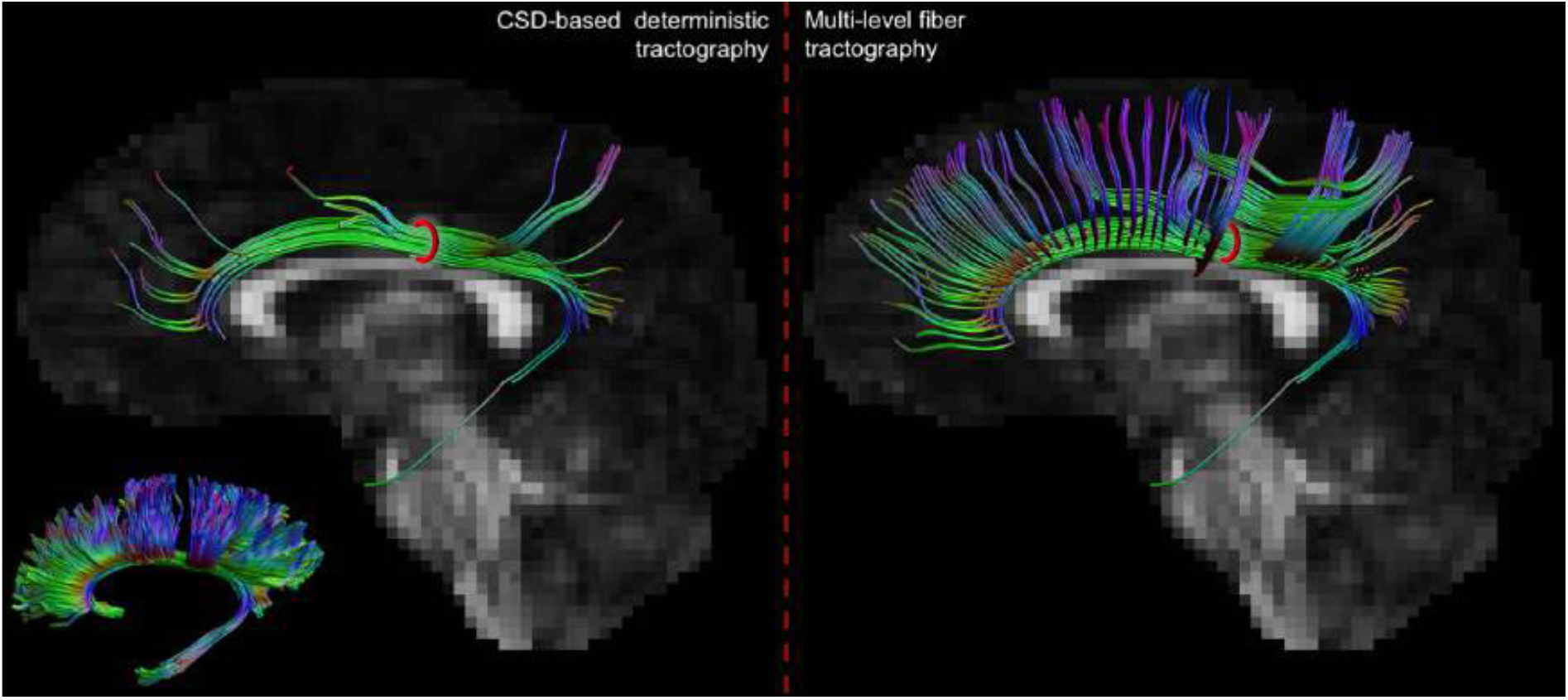
The cingulum bundles reconstructed by the deterministic CSD-based approach (left) and MLFT (right) from the same seed region (red). The cingulum bundle from the ISMRM 2015 challenge ground truth data Maier-Hein et al. (2017) is shown in the bottom left corner for anatomical reference. The delineated pathways correspond better to the physical structure of the cingulum in comparison to the segmentation obtained by deterministic CSD-based tracking. The results are characterized by extensively improved fanning in the anterior and posterior parts of the bundle (Stieltjes et al., 2013; Catani et al., 2012). The seeds were located on the edge of Broadmann areas 23 and 24. The number of seeding points was selected empirically. The target region was chosen based on the White Matter Query Language (Wassermann et al., 2016) cingulum query description adjusted to include only the regions containing the end points of the bundle. Other tracking parameters were set as in the Experiment 3 except for the FOD peak value threshold being set to 0.01.

**Figure S3:**
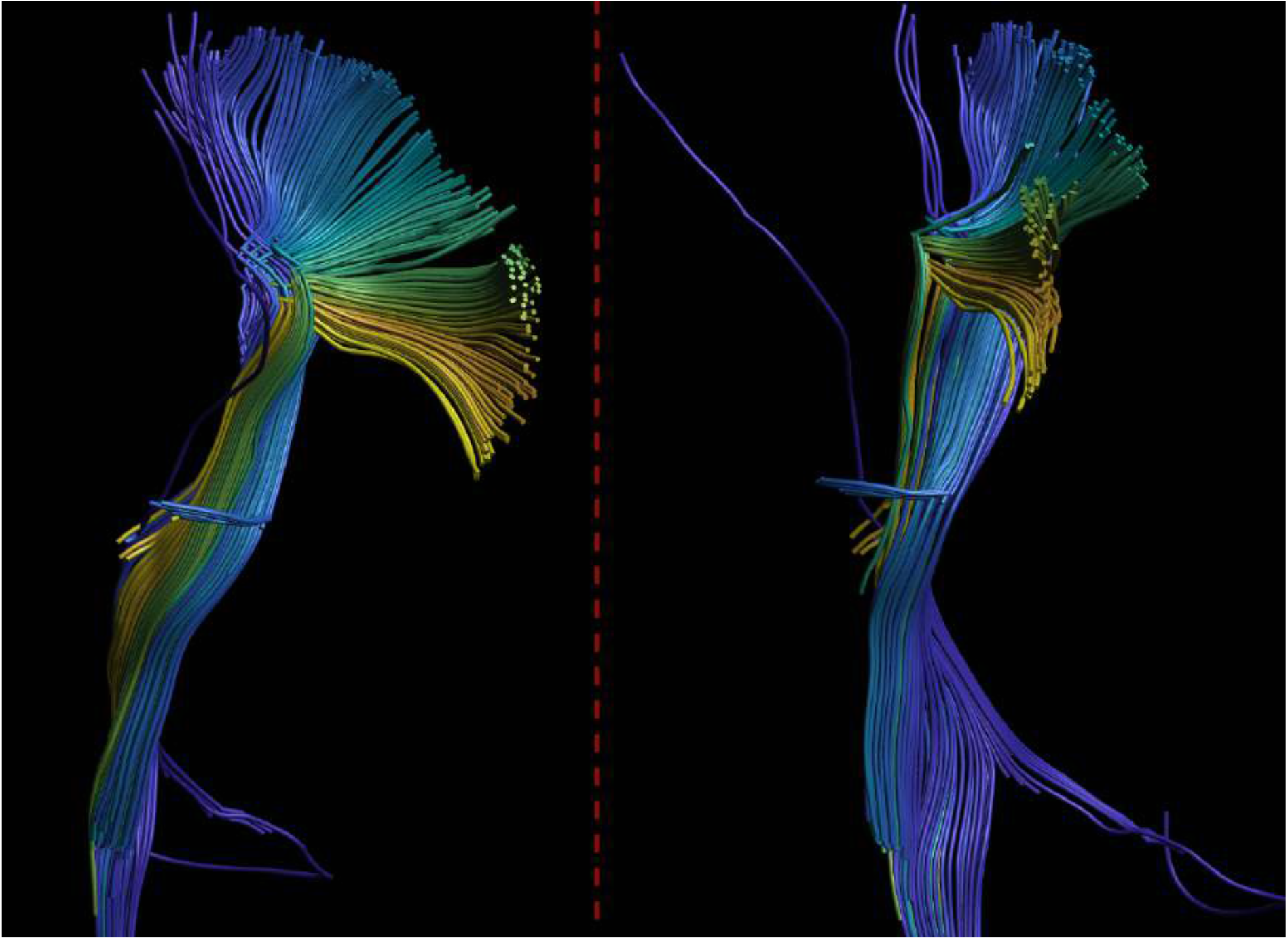
Coronal (left) and sagittal (right) views of the non-filtered left CST reconstructed using the MASSIVE data with pathways colored as in Fig. 17. The non-filtered bundle also maintains a clear topographic organization.

